# Honey wasps differ from other wasps in possessing large gut communities dominated by host-restricted bacteria

**DOI:** 10.1101/2024.08.27.609949

**Authors:** Jo-anne C. Holley, Alexia N. Martin, Anna T. Pham, Jennifer Schlauch, Nancy A. Moran

**Author notes:** Address correspondence to Jo-anne C. Holley.

## Abstract

Honey-feeding social bees, including honey bees, bumbles bees, and stingless bees, possess distinctive gut bacterial communities that provide benefits to hosts, such as defense against pathogens and parasites. Members of these communities are transmitted through social interactions within colonies. The Mexican honey wasp (*Brachygastra mellifica)* represents an independent origin of honey-storing within a group of social Hymenoptera. Honey wasps feed on and store honey, but, unlike bees, they prey on other insects as a protein source, and do not consume pollen. We surveyed the gut bacterial communities of Mexican honey wasps across sites within Texas using 16S rRNA profiling, and we estimated bacterial titer per bee using qPCR. For comparison, we also surveyed non-honey feeding wasps from six families, collected in the same region. We found that honey wasp communities are dominated by characteristic bacterial species. In contrast, other wasps had lower absolute titers and more variable communities, dominated by environmental bacteria. Honey wasps from all sampled nests contained strains of *Bifidobacterium* and *Bombilactobacillus* that were closely related to symbionts of bumble bees and other bees, suggesting acquisition via host-switching. Some individuals also harbored a close relative of *Candidatus* Schmidhempelia bombi (Orbaceae), an uncultured bumble bee symbiont, again suggesting host-switching. The most prevalent species was an uncultured *Lactobacillus* that potentially represents an independent acquisition of environmental *Lactobacillus*. The transition to honey feeding, combined with a highly social life history, appears to have facilitated the establishment of a bacterial community with similarities to those of social bees.

**IMPORTANCE:** Honey-feeding social insects such as honey bees and bumble bees have conserved gut bacterial communities that are transmitted among nestmates. These bacteria benefit hosts by providing defense against pathogens, and potentially by contributing to pollen digestion. The bacterial communities of wasps are less studied. Whereas most wasps are carnivorous and consume nectar, honey wasps (*Brachygastra* spp.) store and eat honey. Here, we address the consequences of this dietary shift for the gut community. Using field collections of Mexican honey wasps and other co-occurring wasps, we found that honey wasps have distinctive gut bacterial communities. These include several bacteria most closely related to bacteria in bumble bees, suggesting their acquisition via host-switching. Solitary wasps and social wasps that do not make honey have smaller gut communities dominated by environmental bacteria, suggesting that honey feeding has shaped the gut bacterial communities of honey wasps.

## INTRODUCTION

Gut microbial communities often provide numerous benefits to insect hosts (1). They can protect against pathogens or parasites (2–4) enhance or become essential for physiological development (5), metabolize indigestible substrates consumed by the host (6, 7) or neutralize toxic compounds (8, 9). Depending on the host species, microbial community members can be highly host-specific or can consist of bacteria widespread in the environment. Well-studied examples of host-restricted bacterial communities are found in the honey-feeding bees, which belong to a clade consisting of honey bees (genus *Apis*), bumble bees (genus *Bombus*) and stingless bees (tribe Meliponini). These social bees are associated with a bacterial community that is relatively consistent among different species, reflecting a shared evolutionary history (10). Most species harbor representatives of five bacterial lineages that appear to have colonized a common ancestor 80 million years ago and to have subsequently diversified with hosts, though a few bee lineages have lost ancestral associations (11). For example, in the best-studied host, the western honey bee (*Apis mellifera*), adult guts are dominated by these five bacterial groups (sometimes called the “core microbiome”), with several other “accessory” bacterial species present more erratically. These bacteria have been implicated in digestion and detoxification of the nectar, honey, and pollen diet of their hosts (8, 12–16). Conversely, solitary bees (such as non-corbiculate bees) have more variable gut communities containing bacteria commonly associated with diverse bee lineages, flowers (nectar and pollen), and other environmental sources (17–21). Bee gut microbial communities can vary with pathogens (22) and urbanization (23, 24), although some of these bacteria may also protect against pathogens (25).

Dietary shifts are frequently associated with a corresponding shift in gut microbial communities (26, 27). A new diet may facilitate or necessitate the acquisition of new symbionts including symbionts that can contribute to nutritional processing. For example, omnivorous, litter-feeding cockroaches have similar gut bacteria regardless of diet composition (28); however, the switch to wood-feeding in *Cryptocercus* cockroaches was accompanied by the acquisition of lignocellulose-degrading protists (29). These symbionts are maintained in the wood-feeding lower termites but were later lost in the soil and litter-feeding higher termites, which possess polysaccharide-degrading fungal and bacterial symbionts that resemble the microbial communities of litter-feeding cockroaches (7, 30–32). On a shorter evolutionary timeline, the vulture bee, *Trigona necrophaga*, switched from pollen and honey to an obligately necrophagous diet. This diet shift is linked to a higher abundance of environmentally acquired acidophiles and more variable bacterial communities compared to related honey-feeding bees (33).

Routes of transmission can also affect microbial community composition. Social and gregarious insects can transfer symbionts among group members by direct contact (34), as documented for honey bees (35) and other groups (36). Sociality, however, is not essential for maintaining a conserved, host-restricted microbial community. *Xylocopa* carpenter bees have conserved bacterial communities but limited social interactions within communal nests (37, 38). Brood provisions serve as a microbial transmission route for some solitary bees (21, 39). However, the gut communities of many solitary bees are acquired from environmental sources (pollen and nectar) and are thus heterogeneous (20).

Here, we investigate the gut bacterial communities of a honey-feeding wasp, representing an independent evolution of honey-feeding. Our focal species is the Mexican honey wasp, *Brachygastra mellifica*. As for honey bees, their colonies contain thousands of individuals, can live for several years, and new nests are established by swarming (40, 41). The honey wasps (*Brachygastra* spp., Vespidae paper wasps) store and feed on honey rather than nectar, as documented in three species (*B. mellifica*, *Brachygastra lecheguana*, and *Brachygastra scutellaris*) (41, 42). This diet shift is rare among wasps; only *Polistes annularis* is known to store honey to overwinter (43), although *Mischocyttarus* spp. store honey short-term (44). To address how this shift may have affected bacterial community composition, we characterized the gut bacteria of Mexican honey wasps across their range in Texas (Fig. 1). We sequenced the V4 region of the 16S rRNA gene to determine the bacterial community composition of 63 honey wasps from 13 nests. Honey wasp gut bacterial communities were compared to those of social *Polistes* paper wasps and several solitary wasps. Our findings indicate that honey wasps acquired a unique gut community dominated by bacterial lineages most closely related to bacteria associated with honey-feeding bees, plus an uncultured species of *Lactobacillus*. This composition differs from that observed for other sympatric wasps, including social paper wasps (genus *Polistes,* Vespidae) and solitary wasps, both of which have more variable gut bacterial communities consisting of bacteria found in other environments.

**Figure 1:**
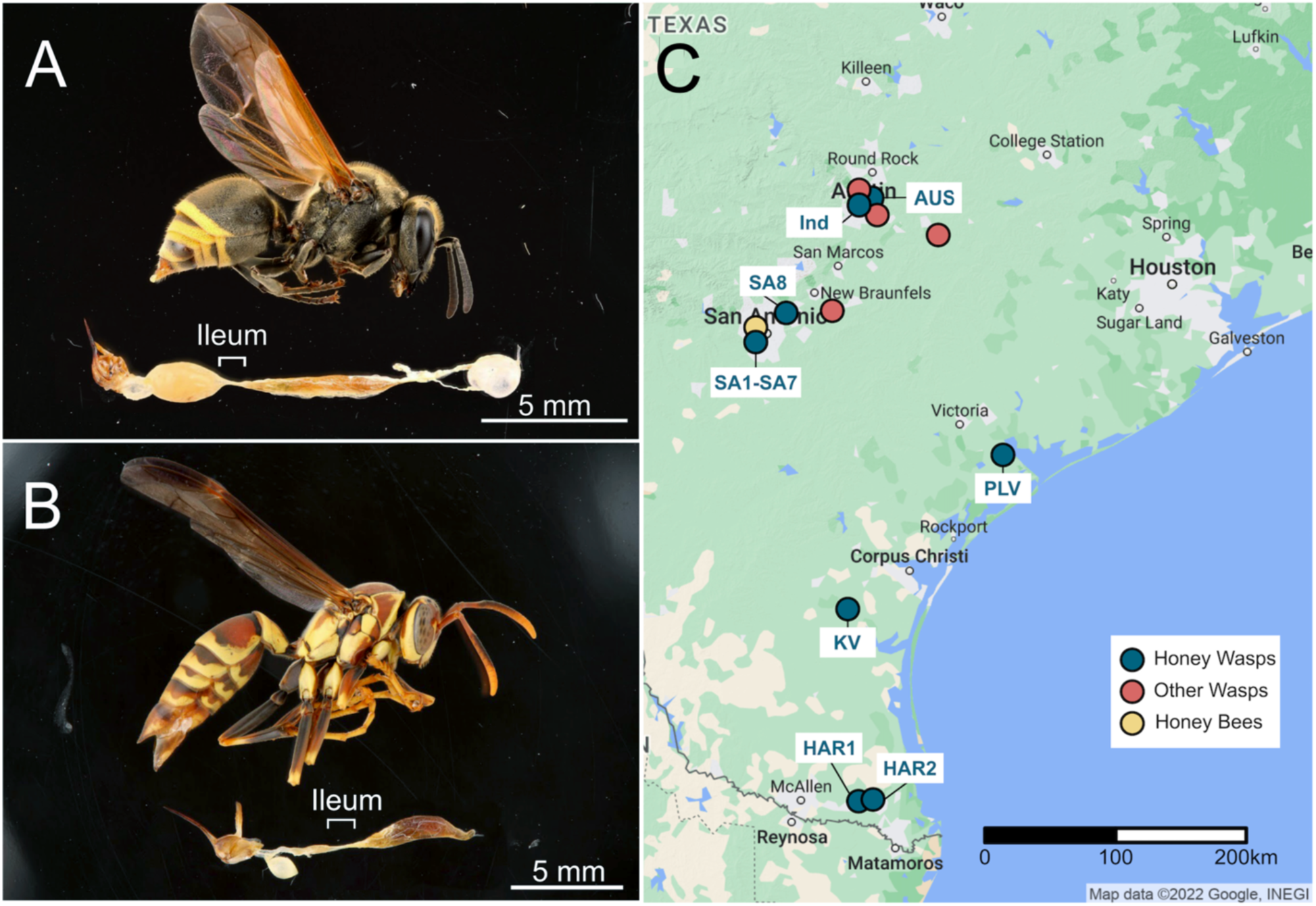
Specimen images and collecting locations. (A) Mexican honey wasp, *Brachygastra mellifica*, (B) common paper wasp, *Polistes exclamans*. Specimens have their guts removed to show the internal structures of the midgut, ileum, and hindgut. The bottom section of the images were cut and rotated to orientate the midguts to the right and sting to the left. (C) Map of Texas showing the collecting sites of honey wasps, *Polistes* paper wasps and solitary wasps, and honey bees.

## RESULTS

### Gut bacterial communities of honey wasps

We sampled 63 Mexican honey wasps, including three to six workers sampled from 13 nests, and five workers collected from flowers in Austin (Fig. 1). For comparison, we sampled 47 other wasps, but only 32 of these wasp samples (15 social *Polistes* paper wasps and 17 solitary wasps) had adequate read counts to be included in the analysis (Table S1-S2, explanation of process in Supplementary materials and methods). One honey wasp sample was also removed, due to low read numbers after filtering (Table S1-S2). Three blank samples were included as negative controls to detect contamination during DNA extractions and sequencing. This resulted in 13 - 589 reads from 11 ASVs after filtering (Table S2).

*Bifidobacterium* and *Lactobacillus* (Fig. 2) were present in all honey wasp nests; however, some individuals did not possess these bacteria. Seven *Lactobacillus* ASVs (2, 7, 8, 25, 53, 62, and 72) and three *Bifidobacterium* ASVs (1, 3, and 5) were mostly confined to honey wasps, although the *Lactobacillus* ASV2 was present in very low abundance in four solitary wasps. Honey wasps also possessed *Convivina* (ASV17 and ASV26), *Fructilactobacillus* (ASV13), and *Bombilactobacillus* (ASV21) throughout the Texas range, but these were not present in all individuals (Fig. 2B). Honey wasps from some San Antonio nests had bacteria related to *Ca.* Schmidhempelia bombi (ASV14 and ASV30), an undescribed Neisseriaceae (ASV27), and *Asaia* (ASV36). The honey wasps collected in Austin, from a nest (AMP in Fig. 1) or from flowers, had more variable bacterial communities and were not dominated by *Lactobacillus* and *Bifidobacterium*. Some AMP samples had bacterial communities dominated by *Fructilactobacillus*, while others had high abundance of *Ca*. Stammerula and *Convivina*. Several Austin honey wasps possessed two Enterobacteriaceae (ASV4 and ASV6) also found in solitary wasps.

**Figure 2:**
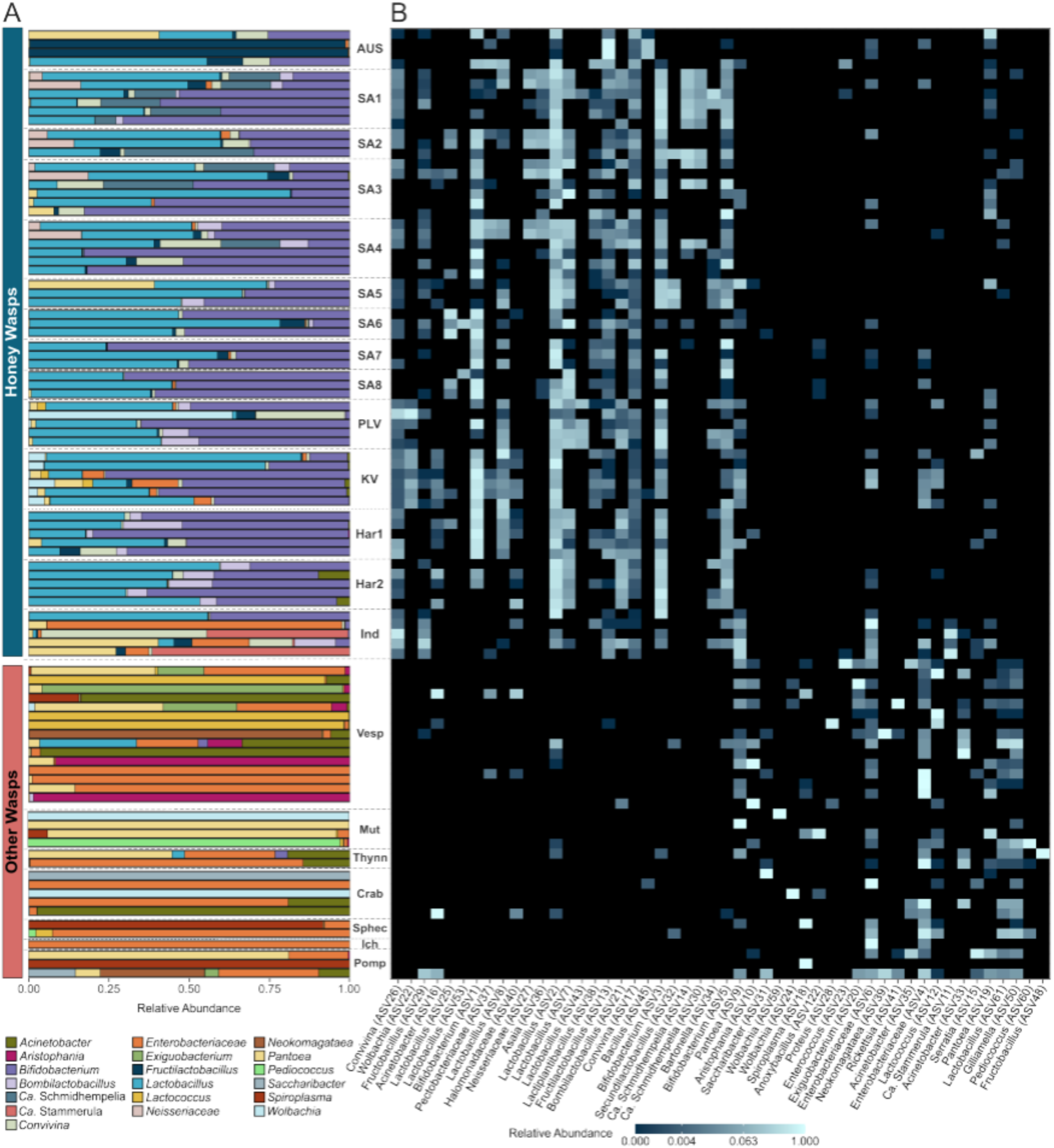
Relative abundance of (A) the top 16 bacteria genera and (B) the top 50 amplicon sequence variants (ASVs) from honey wasps, other wasps, and honey bees. Honey wasps are grouped by nest, except five individuals collected in Austin (Ind) based on the rarefied data. Nests are named based on their location and organized on the plot from north (top) to south Texas (bottom). Other wasps are grouped by family (Vesp = Vespidae, Mut = Mutillidae, Thynn = Thynnidae, Crab = Crabronidae, Sphec = Sphecidae, Ich = Ichneumonidae, Pomp = Pompillidae). Honey wasps typically possess a characteristic bacterial community that distinguishes them from other wasp species.

Based on the “core microbiome” analysis, some ASVs of *Lactobacillus*, *Bifidobacterium*, *Bombilactobacilus*, *Fructilactobacillus*, and *Convivina* occur in most honey wasp individuals in high abundance (Fig. 3). An NMDS ordination of the Bray-Curtis dissimilarity index shows that the communities of honey wasps cluster together apart from those of other wasps (Fig. 4A, Table S5, PERMANOVA, F = 15.968, p = 0.001). Beta dispersions significantly differed among the honey wasps and the combined wasps (Table S6, betadisper, F = 34.933, p = 0.001).

**Fig. 3.**
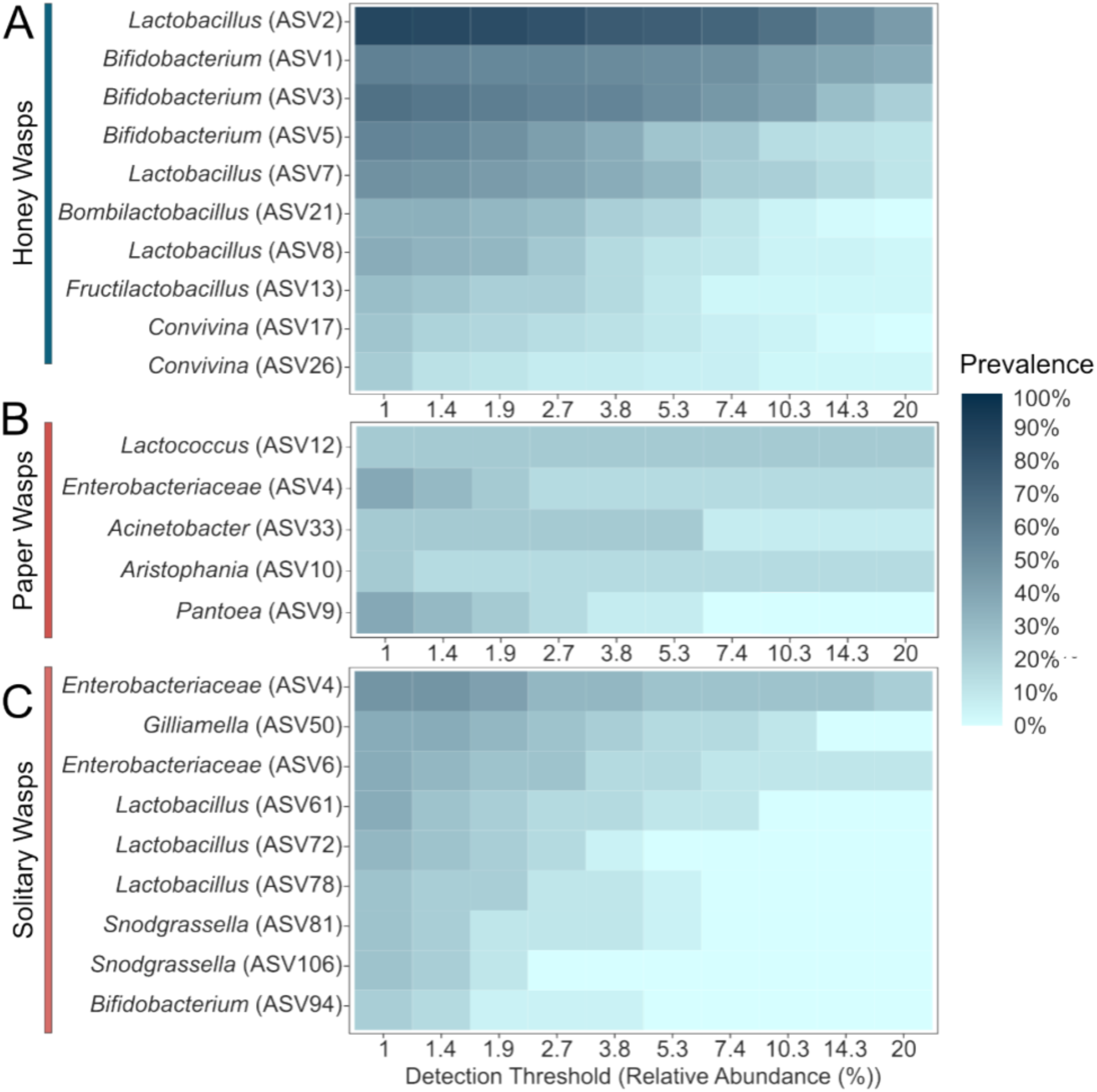
“Core microbiome” analysis of the rarefied data set showing three insect groups (A) honey wasps, (B) *Polistes* paper wasps, and (C) solitary wasps studied. Prevalence is the proportion of insect samples in each group containing ASVs at detection thresholds ranging from 1% to 20% of the total read abundance. Many solitary wasps are excluded due to lack of amplification, and some included samples had low titers, raising the possibility that contaminants or dead bacteria in food dominate.

**Figure 4:**
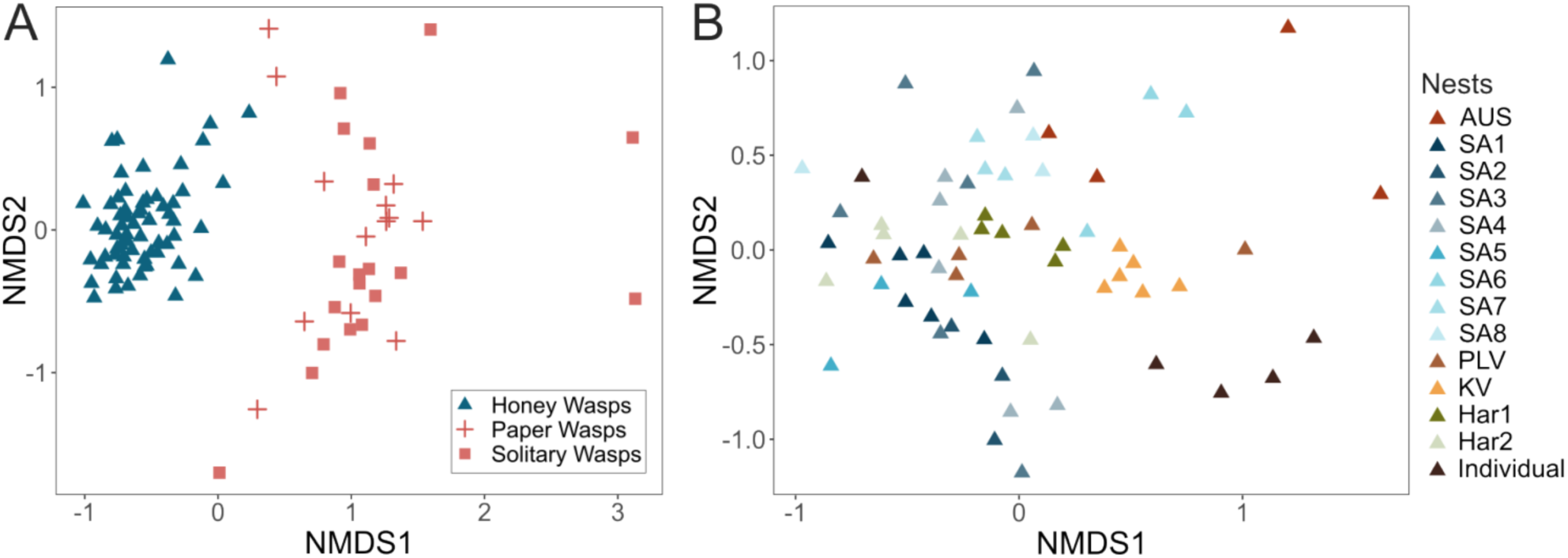
Nonmetric multidimensional scaling (NMDS) ordination of Bray-Curtis dissimilarity index of the rarefied data set. (A) Honey wasp gut bacterial communities compared to those of *Polistes* paper wasp and solitary wasps (stress = 0.19). (B) Honey wasp gut bacterial communities categorized by nest of origin (stress = 0.24).

### Bacterial communities of *Polistes*, solitary wasps

The gut bacterial communities of *Polistes* and solitary wasps sometimes contained *Bifidobacterium* (ASV94) or *Lactobacillus* (ASVs 61, 72, 111). Still, these ASVs were absent from honey wasps (Fig. 2B). *Polistes* and solitary wasps shared Enterobacteraceae ASVs (ASV4 and ASV6) with honey wasps. They more often contained *Aristophania*, *Lactococcus*, *Acinetobacter*, and *Exiguobacterium*, *Pantoea*, and *Spiroplasma* (Fig. 2). “Core microbiome” analysis showed that *Polistes* were typified by the presence of the these bacteria(Fig. 3), but no bacteria were detectable in more than 40% of *Polistes* individuals sampled, indicating that *Polistes* bacterial communities are often absent or tiny. The *Polistes* shared some ASVs with the 18 solitary wasps (Enterobacteriaceae ASV4, *Pantoea* ASV9 and ASV19, *Lactobacillus* ASV61 and ASV72), Enterbacteriaceae (ASV6) had higher relative abundance in the solitary wasp samples compared to *Polistes* samples. Solitary wasp bacterial communities were highly variable and dominated by bacteria such as *Acinetobacter*, *Pantoea*, *Pediococcus*, *Wolbachia*, and *Spiroplasma* (Fig. 2aA). “Core microbiome” analysis indicated the most prevalent bacteria were *Gilliamella* (ASV50), three *Lactobacillus* ASVs (ASVs 61, 72, and 78), and two *Snodgrassella* (ASV81 and ASV106) (Fig. 3). These bacteria were all in low abundance (fewer than 12% of total reads in samples) and are not represented among the ASVs in Fig.2A. Notably, the ten most abundant ASVs in honey wasps were not among the most abundant ASVs in *Polistes* and other wasps.

### Absolute titers of honey wasp gut bacterial communities

Based on qPCR of a region of the bacterial 16S rRNA gene, honey wasps have substantial gut communities, averaging greater than 10^8^ gene copies per wasp individual, with relatively little variability in community size (Fig. 5). In contrast, the absolute titers of gut bacteria in *Polistes* and in solitary wasps were lower and highly variable, ranging from below the limit of detection to 10^10^ copies of the bacterial 16S rRNA gene (Fig. 5, Table S8). *Polistes* wasps had a median of 3.1 x 10^5^ copies of 16S rRNA gene, ranging from 10^3^ to 10^7^ copies, and solitary wasps had a similar wide range of bacterial load (Table S9, Dunn’s test, z = −1.662, adjusted p = 0.290).

**Figure 5:**
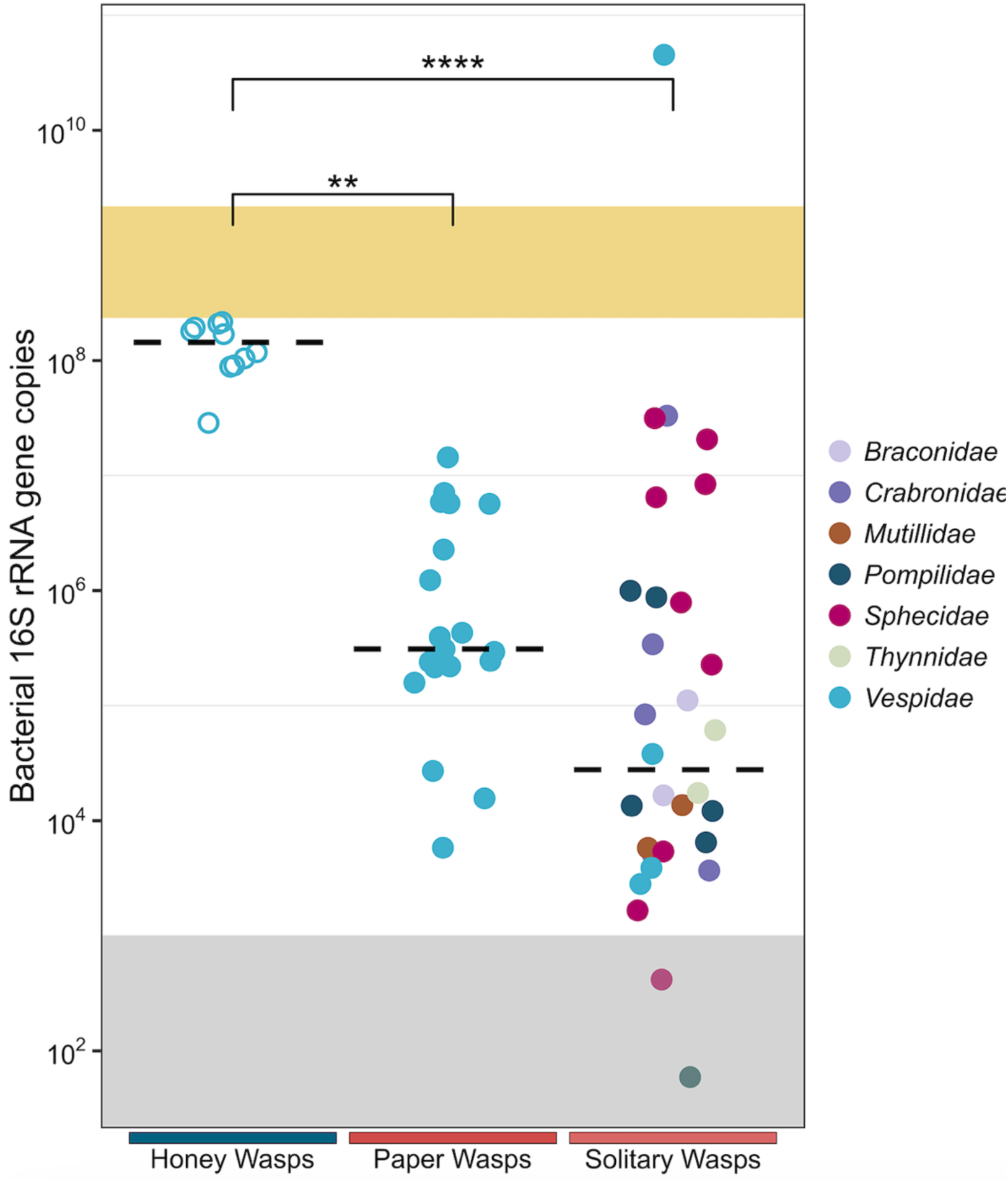
Bacterial 16S rRNA gene copies per individual host/host gut for honey wasps, social *Polistes* paper wasps, and solitary wasps. Dashed line is the median. The gray shaded area is the region below the detection threshold, and the yellow shaded area represents the range of 16S rRNA gene copies for *Apis mellifera* from Kwong et al. (10). Honey wasps (n = 10) were from nests in San Antonio (SA1 and SA3), *Polistes* samples represent five species (n = 19), and solitary wasps represent 7 families (n = 30, including four solitary eumenine vespid wasps). Honey wasp gut bacterial communities significantly differed from those of *Polistes* paper wasps and solitary wasps in all pairwise comparisons (honey wasps:paper wasps, z = −3.417, adjusted p = 0.002; honey wasps:solitary wasps, z = −4.990, adjusted p = 1.8 x 10^−6^).

### Bacterial community diversity among honey wasp nests

Honey wasp bacterial composition varied among sample locations and varied significantly among nests (Fig. 4B, Table S5 and S7, PERMANOVA, F = 2.9927, p = 0.001; betadisper, F = 1.2641, p = 0.253). No nests had distinctly different gut communities based on pairwise comparisons of honey wasps from all 13 nests (Table S7, adjusted for multiple comparisons). *Bifidobacterium* and *Lactobacillus* were the primary members of the bacterial communities and present in all nests, but there were multiple ASVs for each that were differentially distributed among nests.

Honey wasp *Bifidobacterium* was represented by three ASVs. Indicator species analysis revealed that ASV1 was significantly associated with five nests while ASV3 was associated with four different nests (Table S10). *Bifidobacterium* ASV1 and ASV3 rarely co-occurred in the same host, but ASV3 typically occurred with *Bifidobacterium* ASV5 (Fig. 2B).

*Lactobacillus* ASV2 was present in all nests, but the indicator species analysis associated ASV2 with five particular nests (Table S10, indicator value = 0.602, p = 0.022). Other *Lactobacillus* ASVs were associated with a subset of nests. For example, *Lactobacillus* ASV25 was associated with San Antonio nest 6 and ASV43 with the Port Lavaca nest (Table S10; ASV25: indicator value = 0.819, p = 0.03; ASV44: indicator value = 0.869, p = 0.0022).

Some bacteria had a limited association with only a few nests. For example, San Antonio nests 1, 2, and 3 were characterized by the presence of ASVs clustering with *Ca.* Schmidhempelia (Table S10; ASV14: indicator value = 0.639, p = 0.0418; ASV30: indicator value = 0.636, p = 0.0439). Additionally, the Austin nest showed high abundance of *Fructilactobacillus* (Fig. 2B, Table S3, Table S10:indicator value = 0.715, p = 0.0412).

### Laboratory isolation of honey wasp bacteria

To verify that the ASVs from honey wasps represented living bacteria, we isolated pure cultures of some community members. These included a member of the *Ca.* Schmidhempelia cluster (wjB12) and several strains of *Bifidobacterium* (efB7, efB4, wjB1) from two San Antonio nests (SA1 and SA7) grown on heart infusion agar (HIA) with 5% sheep’s blood under 5% CO_2_ at 35°C. A *Fructilactobacillus vespulae* isolate (sjhB6) and an additional *Bifidobacterium* isolate (sjhB46) were also cultured in MRS broth under anaerobic conditions at 31°C. The identities of these cultures were verified with 16S rRNA gene sequencing.

### Phylogenetic relationships of key honey wasp bacteria

We constructed phylogenies for three bacterial groups prevalent in honey wasp bacterial communities belonging to Orbaceae, *Bifidobacteria*, and Lactobacillaceae. Within Orbaceae, the honey wasps possessed two ASVs and a bacterial isolate (strain wjB12) that form a well-supported clade together with *Candidatus* Schmidhempelia bombi, an uncultured Orbaceae known from bumble bees and carpenter bees (Fig. 6A) (37, 45, 46).

**Figure 6:**
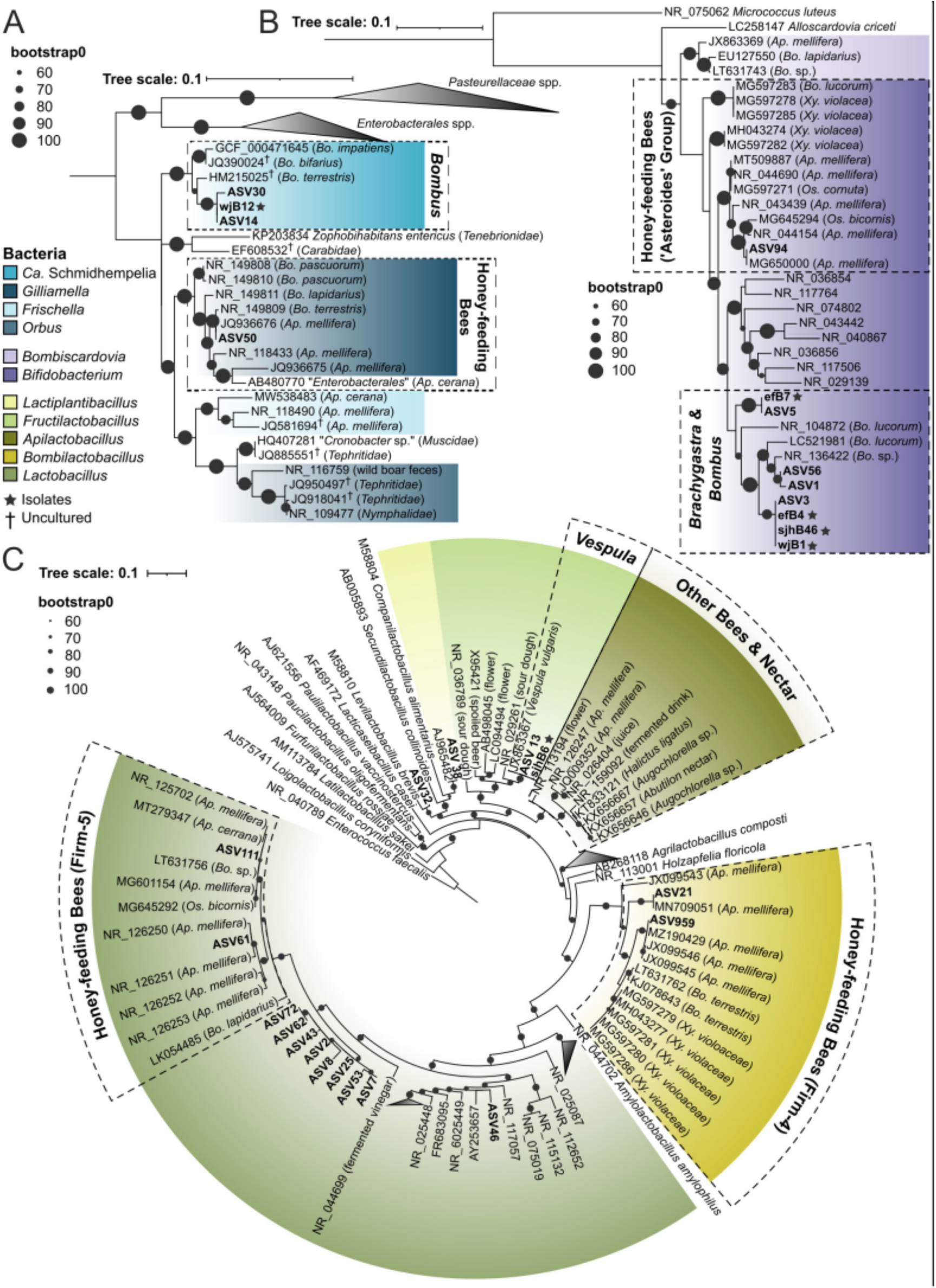
Phylogenies of the 16S rRNA gene of the major bacterial taxa from the honey wasps. (A) Orbaceae, (B) Bifidobacteriaceae, and (C) Lactobacillaceae. Phylogenies were inferred by maximum likelihood, with bootstrap support indicated by black circles. Colors represent bacterial genera or clades associated with bees, host source listed in parentheses. ASVs from this study are highlighted, and the asterisks denote bacteria isolated from honey wasps.

Three *Bifidobacterium* ASVs and numerous *Bifidobacterium* isolates from honey wasps form a clade with multiple species of *Bifidobacterium* isolated from several species of European bumble bees (*B. commune*, *B. bohemicum*, *B. bombi*) (Fig. 6B). This clade is not closely related to the *Bifidobacterium ‘asteroides’* clade associated with honey bees, bumble bees, carpenter bees, and *Osmia* mason bees (10, 37, 55).

Honey wasps also possess a variety of bacteria in the family Lactobacillaceae. Eight near-identical ASVs represent an undescribed species of *Lactobacillus* that forms a sister group to the clade of *Lactobacillus* associated with honey bees, bumble bees, stingless bees, and *Osmia* bees (containing *Lactobacillus melliventris* and formerly referred to as “Firm-5”) (Fig. 6C). Seven of these *Lactobacillus* ASVs were found primarily in honey wasps, while ASV72 was found in *Polistes* paper wasps. Representatives of this ASV cluster remain uncultured. The 16S rRNA gene amplicon variation suggests there is strain diversity within the honey wasps with ASV2 found in 92% of honey wasps sampled.

Another Lactobacillaceae group, *Bombilactobacillus* (containing *Bombilactobacillus mellifer* and formerly called “Firm-4”), was sequenced from multiple honey wasps but remains uncultured (Fig. 6C). Previously the genus was known only from honey bees, bumble bees, stingless bees, and carpenter bees (10, 11, 37, 47). The sequences from honey wasps are close to those from *Bombilactobacillus apium* isolated from the Asian honey bee, *Apis cerana*. Additionally, honey wasps possess the vespid wasp-associated *Fructilactobacillus vespulae*, a *Lactiplantilactobacillus*, and another Lactobacillaceae (Fig. 6C).

## DISCUSSION

### Major findings

Honey wasps represent an independent evolution of honey storage and feeding in social Hymenoptera, and we explored whether this dramatic shift in diet was linked to changes in the gut bacterial community. We found that honey wasps have bacterial communities that differ from those of other wasps, both social and solitary, but that exhibit some similarities with communities of social bees (Figs 2A, 2B). First, as in the social bees, honey wasps have large bacterial communities with relatively consistent absolute gut titers, as compared to other wasps, which have bacterial titers that are over two orders of magnitude smaller on average and that are highly variable in size (Fig. 5). As estimated by qPCR, sizes of honey wasp bacterial communities are comparable to those of honey bee species, at about 10^8^ copies of rRNA genes per bee (10). Second, honey wasp bacterial communities have relatively consistent composition dominated by lineages that are host-restricted, as in social bees. Third, certain members of the honey wasp bacterial communities are most closely related to taxa in microbiomes of social bees (Fig. 6). However, other bacteria characteristic of honey wasp gut communites are not linked to bacteria from social bees.

### Gut bacterial communities of honey wasps

Honey wasps have a diverse consortium of Lactobacillaceae that is distinct from that of honey bees or the other wasps included in this study. All honey wasps host an uncultured *Lactobacillus* which belongs to a clade of honey bee-associated *Lactobacillus* near *L. melliventris* (formally Firm-5); multiple ASVs were detected indicating strain diversity in this taxon (Fig. 6C). Despite many efforts, this *Lactobacillus* remains uncultured.

*Bombilactobacillus* ASVs in honey wasps are closely related to *B. apium* isolated from *Apis cerana* (Fig 6C) (47). Honey wasps also consistently possess three related *Bifidobacterium* ASVs that belong to the same clade as *Bifidobacterium* species (*commune*, *bombi*, *bohemicum*) isolated from European bumble bees (Fig. 6B) (48, 49). We successfully cultured two strains (corresponding to ASV3 and ASV5) from honey wasp guts, demonstrating that these ASVs represent living bacteria. Individual wasps had either ASV1 or ASV3 in high abundance, not both, suggesting competition or another incompatibility.

Though honey wasp bacterial communities had relatively consistent composition, some bacteria were erratically present. Several nests of honey wasps in San Antonio had ASVs closely related to *Ca.* Schmidhempelia bombi, known from metagenomic samples of *Bombus* impatiens (Fig. 6A)(45). The ASVs from honey wasps nested, with strong support, within a clade previously known only from bumble bees (Fig. 6A). Honey bees, stingless bees, and bumble bees share gut bacteria, including *Lactobacillus*, *Bifidobacterium*, and *Bombilactolacillus* that were acquired by a common ancestor approximately 80 million years ago (10), and the bumble bee associate *Ca*. Schmidhempelia was acquired by bumble bees later (50). Associations between honey-feeding bees and their bacterial symbionts predate the evolution of the genus *Brachygastra*, which arose approximately 13.6 million years ago (51). Mexican honey wasps and stingless bees have co-occurred in South, Central and southern parts of North America during that time, and *Bombus* arrived in the Neotropics several million years ago (52), providing ample time for the lineages to interact. The shared food source of nectar may have enabled transfer between honey wasps and honey-feeding bees.

Also present in the honey wasps were bacteria commonly associated with other, non-honey-feeding bees. These include *Lactococcus, Convivina*, and some *Lactobacillaceae* such as *Lactiplantibacillus*. Three honey wasps collected individually from Austin had high abundances of ASVs related to *Candidatus* Stammerula and with 99% sequence identity to an uncultured symbiont of tephritid flies (53, 54).

While honey wasps appear to have acquired some bacterial lineages found in honey-feeding bees, they lacked members of the *Bifidobacterium “asteroides”*, *Snodgrassella,* or *Gilliamella* clades, which are found in bumble bees, honey bees and many stingless bees (10, 55). Honey wasps also lack the “Firm-5” *Lactobacillus* species (*L. melliventris* group). associated with social bees but possess an undescribed relative. *Snodgrassella* and *Gilliamella* form a biofilm along the ileum of honey bees and bumble bees (12, 50). Also missing are the “accessory” bacteria that are occasionally sampled in honey bee bacterial communities, such as species in the genera *Apilactobacillus*, *Bombella*, *Arsenophonus*, *Bartonella*, *Parasaccharabacter*, *Apibacter*, and *Frischella perrara* (56, 57).

Our study on honey wasps adds to others suggesting that some vespid wasps have reliable associations with gut bacteria. The subfamily Vespinae contains four genera, of which two have been the focus of research on bacterial associates. *Vespula pensylvanica*, *Vespula vulgaris* and *Vespa velutina* are all associated with several bacteria that we found in honey wasps. *Vespa velutina* nests consistently possess *Lactobacillus* and *Bifidobacterium* in all life stages (58), as we found for honey wasps. *Fructilactobacillus vespulae*, which we found in most honey wasp nests, has been isolated or sequenced from *Vespula vulgaris* in China, *Vespula pensylvanica* in the US (59), and *Vespa velutina* in Belgium. We detected *Convivina* in some individuals from all sampled honey wasp nests, but not in other wasps. *Convivina intestini* has been isolated from *Bombus terrestris* (60) and *Convivina praedatoris* was found in a single *V. velutina* nest collected in Belgium (outside of the Korean native range) (61). Similarly, we detected *Lactiplantibacillus*, another possible member of the *V. velutina* community, in 43% of sampled honey wasps. In a study of bacterial communities of five *Vespa* species in their South Korean native range, *Convivina* and *Lactiplantibacillus* were not among the top 10 bacterial genera (62), so these associations may be site-specific.

### Gut bacterial communities of other wasps

We found that social *Polistes* paper wasps and solitary wasps have relatively small gut communities that vary in composition and absolute bacterial titers. Their bacterial titers, estimated as copies of 16S rRNA operons, were 10^4^-10^6^, far lower than for honey wasps or honey bees (Fig. 5). These values likely overestimate average community sizes, as almost half of these wasp samples failed to produce sufficient 16S rRNA amplicons and were excluded. In contrast, all samples for honey wasps produced sufficient amplicons. Also, we note that community profiles obtained from PCR amplicons include reads from bacteria present in the food and from extracellular DNA from dead bacteria, in addition to bacteria colonizing the gut. The possibility that these other sources dominate profiles is especially great for low bacterial titer communities consisting of environmental bacteria, such as those in the *Polistes* and solitary wasps. Since many species of bees and wasps can visit the same flowers, where they may contaminate nectar, live or dead cells from other host species may be ingested in food.

The *Polistes* paper wasps and solitary wasps had highly variable gut community compositions, often dominated by environmental bacteria, including environmental Lactobacillaceae. They lacked all major members of the honey wasp bacterial community. Some ASVs erratically present in honey wasps were shared with *Polistes* and solitary wasps, including some from Enterobacteriacae, *Pseudomonas*, *Lactococcus*, and some others found in environmental sources such as nectar and other Hymenoptera (20, 37, 56, 59). However, these had low prevalence and low abundance in honey wasps (Fig. 4, Fig. 6).

In the four species of social *Polistes,* sequences corresponding to *Aristophania*, *Gilliamella*, *Acinetobacter*, and *Lactobacillus* were found erratically (Fig. 2A). These have been previously detected in flower nectar, solitary bees (17, 63, 64), *Vespa* spp. (62), and *Vespula pensylvanica* (59). We found *Gilliamella* sp. (ASV 50) in a variety of wasps including species from Mutillidae, Crabronidae, Sphecidae, and *Polistes* in low abundance (Table S3). *Gilliamella* species are among the bacteria consistently present in honey bees and bumble bees (58), and sequences from some strains have been detected in native bee species and in nectar (65). These are likely to represent DNA present in ingested nectar, as honey bees are the most common visitors to flowers at all sites.

*Polistes* workers also feed on nectar and share it with larvae, as do many solitary wasps, explaining how they acquire some bacteria. However, many adult bees have bacterial communities similar to those in their pollen provisions (20, 21, 23, 24, 63, 65), whereas *Polistes* derive protein from insect prey and do not store pollen. *Polistes* are social and thus have a means of bacterial transmission among nestmates and generations, yet their gut communities are composed of environmental bacteria found in nectar and in solitary bees. It appears that both *Polistes* and solitary wasps acquire bacteria from environmental sources based on the wasps sequenced in this study. Our findings suggest that many wasps, both social and solitary, do not rely on a specific bacterial community, as is the case for some other insects (1, 66).

### Geographic variation in bacterial communities

Mexican honey wasps occur from central Texas south to Panama, so our collection range represents a small part of the *B. mellifica* range. Furthermore, the Mexican honey wasp northern range limit was near the USA/Mexico border until around 1995 (41) and has expanded approximately 500 kilometers north over 3 years. We collected 13 nests in Texas and found significant variation in community composition among nests. Bacterial communities are more similar among nestmates. Differences between nests are due to the presence and abundance of particular ASVs, reflecting strain variation among the major microbiome members. Examples include the uncultured *Lactobacillus*, represented by eight ASVs that form a clade (Fig. 6C), and the *Bifidobacterium* with 4 ASVs that varied among hosts.

All honey wasp individuals had bacterial communities dominated by *Lactobacillus* and *Bifidobacterium*. The exceptions were wasps from the Austin nest, where three of the five individuals were dominated by *Fructilactobacillus vespulae*. Among the five honey wasps that were collected from flowers in Austin, one had the characteristic bacterial profile, and the four others had a higher abundance of Enterobacteriaceae, *Ca*. Stammerula, and *Conviva*. This variation suggests that honey wasps sometimes lose the conserved, host-restricted set of bacteria, which is replaced by a more variable community containing environmental strains, a pattern observed repeatedly in bumble bees (57). These two Austin collection sites sit close to the northernmost extent of the Mexican honey wasp range, suggesting a change in microbial diversity associated with edge populations (though our sampling is not sufficient for conclusions). Two species of English butterflies have microbiomes with decreased diversity and a shift in composition in the northern part of their range (67). Similarly, the microbial communities of range-expanding plants were more similar to each other than to those of conspecifics in the ancestral range (70). Further research into honey wasps throughout Mexico and Central America would reveal whether the bacterial community varies across its range.

### Honey wasp diet and bacterial community

The transition to long-term honey-feeding and storage may have expedited the acquisition of bee-associated bacteria as honey wasps feed on a sugar-rich diet like that of honey-feeding bees. *Bifidobacterium* spp. have extensive abilities to degrade plant-derived polysaccharides (74), and bee-associated *Bifidobacterium* can degrade and use hemicelluloses (13). Honey wasps likely do not ingest significant amounts of polysaccharides, since they do not eat pollen, so metabolic capabilities of their bacterial communities may be very different from those of honey bees and bumble bees, which consume pollen. Investigation of the functional capabilities of the *Bifidobacterium* strains isolated from honey wasps will offer insights into their metabolic capabilities and potential benefits to hosts.

### Conclusions

The Mexican honey wasp gut bacterial community is an amalgamation of bacteria that are related to those from bumble bees or honey bees, from vespine wasps, or from other environmental sources. These contrast with communities from sympatric polistine wasps and solitary wasps, which have erratic bacterial communities composed of environmentally acquired bacteria. The switch from feeding on nectar and insect prey to a stable, year-long supply of stored honey may have facilitated, or been facilitated by, the acquisition of bacteria associated with honey-feeding bees.

## METHODS

### Sample Collection

Honey wasps from 13 nests, individual honey wasps, *Polistes* wasps, and solitary wasps were collected across sites in central and south Texas (Fig. 1, Table S1). Guts were dissected from specimens using sterile technique and stored in 100% ethanol at −80°C. Additional honey wasp guts were preserved in 20% glycerol at −80°C for bacterial isolation.

### DNA Extractions, Sequencing, and qPCR

DNA was extracted from whole guts using the CTAB/phenol-chloroform extraction method (35). The full length or V4 region of the 16S rRNA gene was PCR-amplified and sequenced on an Illumina MiSeq platform at the Genome Sequencing and Analysis Facility (University of Texas at Austin). To estimate total 16S rRNA gene copies per insect as an estimate of bacterial abundance, aliquots of diluted DNA extracts were used as template for qPCR. The observed threshold (CT) cycle values were plotted on a standard curve to determine the absolute copy number.

### Bacterial Isolation and Identification

The preserved honey wasp gut homogenates from two nests (SA1 and SA7) and an individual collected in Austin were grown on HIA + 5% sheep’s blood or in MRS broth at 35°C in 5% CO_2_ or 31°C in anaerobic conditions for 1-3 days. To identify bacterial isolates, the 16S rRNA gene was PCR amplified, and amplicons were sequenced at the DNA Sequencing Facility (University of Texas at Austin).

### Data Analysis and Phylogenetic Reconstruction

Illumina sequencing files were processed in R using the dada2 pipeline to remove poor quality reads and other sequencing artifacts, and to determine unique amplicon sequence variants (ASVs). Taxonomy was assigned using the SILVA 138.2 database (75) and manually confirmed using NCBI BLASTn. Chloroplasts, mitochondria, and unassigned reads were removed, along with contaminants detected in the three blank samples. Read counts were checked after each filtering step; 16 samples were excluded because they 1) lost more than 50% of reads during a post-dada2 filtering or 2) had fewer than 1500 reads after filtering was complete. These issues suggest contamination due to low bacterial biomass (see Supplementary methods, Table S2)(68, 69). Following filtering, the remaining samples had an average of 25,103 reads (range from 2172, to 128,080) (Table S3). To account for uneven sequencing, samples were normalized by rarefying to 95% of the minimum sampling depth, resulting in 2063 reads for each sample (Fig. S2, Table S4). All subsequent analyses were conducted on both the rarefied data set and the full data, and there were no differences in statistical outcomes. The “core microbiomes” of honey wasps, *Polistes*, and solitary wasps were identified using the microbiome package. Indicator species analysis was also performed using the indicspecies package to determine if any ASVs were significantly associated with each honey wasp nest. Statistical differences between gut bacterial communities across honey wasp nests and insect types were analyzed using PERMANOVA (*adonis2*), and assumptions were assessed using beta dispersion analysis. Post-hoc pairwise comparisons with Bonferroni adjustment were performed to directly compare bacterial communities across nests and insect types.

The 16S rRNA gene phylogenies were reconstructed for Lactobacillaceae, Orbaceae, and *Bifidobacterium*, the groups containing most ASVs or isolates from honey wasps. Full length 16S rRNA gene sequences were obtained from GenBank. These were aligned using MAFFT with ASV sequences and full length 16S rRNA from the isolated bacteria. Maximum likelihood phylogenies were reconstructed using IQtree with auto-generated substitution models and 100 standard bootstrap alignments.

## DATA AVAILABILITY

Raw 16S rRNA gene amplicon reads are deposited in the National Center for Biotechnology Information Sequence Read Archive under accession PRJNA932820. ASV sequences, taxonomic assignments, read numbers, metadata can be found in the Supplement. Scripts for the analysis can be found on GitHub (https://github.com/annatpham/honey-wasp-microbiome), and insect specimens are deposited at the University of Texas Insect Collection (UTIC).

## ACKNOWLEDGEMENTS

We thank G. Rankin, H. Koch, S. Bard, D. Fall, S. Biles, J. Hare, M. Reed, B. Freitag, and B. Fehrenkamp for specimen acquisition and collection. Thanks also to A. Santillana for images, A. Wild for specimen ID, and E. Frederick and K. Chow for technical assistance. This work was supported by USDA-NIFA grant (2017-06473), NSF award (NSF 2103208), and the Texas EcoLab program, and undergraduates were supported by the Freshman Research Initiative (UT Austin).

## Notes

### Competing Interest Statement

The authors have declared no competing interest.

### Summary of Updates

The revision has updated analyses and figures.

