## Supplemental materials for "Honey wasps differ from other wasps in possessing large gut communities dominated by host-restricted bacteria"

#### Contents:

Page 2-8: Methods and materials
Page 8-11: References
Fig. S1, page 12: Rarefaction curves
Fig. S2, page 13: Full Lactobacillaceae phylogeny

PAGES 14-21
Table S1: Sample collection information (in an excel file)
Table S2: Reads through the pipeline (excel)
Table S3: ASVs, reads, and taxonomy of the full data set (excel)
Table S4: ASVs, reads, and taxonomy of the rarefied data(excel)
Table S5, page 14: PERMANOVA tests
Table S6, page 14: Homogeneity of dispersion tests
Table S7, page 15-19: Pairwise comparisons
Table S8, page 20: qPCR summary statistics
Table S9, page 20: qPCR comparisons
Table S10, page 21: Indicator species analysis

### Materials and methods

#### Study organisms

The Mexican honey wasp is a polistine paper wasp in the family Vespidae that forms colonies ranging from 3,000 to 20,000 individuals (1, 2). Colonies contain one reproductive female or up to 20% reproductive females, live for an average of three years, and establish by swarming (1, 2). Like honey bees, the wasps feed on honey made from a variety of local nectar sources, and possibly hemipteran honeydew (2), which is stored in the nest in wasp-constructed paper hexagonal structures made of chewed plant fibers and saliva. Honey wasps prey on small soft-bodied insects such as aphids and caterpillars as a source of protein (2, 3). Hunt et al. (4) surveyed pollen of two *B. mellifca* individuals, and they had 2-19 pollen grains in their gut. Our informal observations found little to no pollen in the honey wasp guts. Thus, it is unlikely that pollen serves as a protein source for honey wasps. Among the vespid wasps in general (including *Brachygastra* spp. and *Polistes* spp.), feeding behavior is well documented. Workers forage for insects, these are malaxated (chewed) to extract liquid, and the prey is fed to larvae (4). Nectar is collected and shared among workers, and larvae also supply adults with amino acid-rich saliva (5). *Polistes* spp. engage in bidirectional food-sharing among larvae and adults and trophallaxis among workers (5), although this has not been observed for honey wasps.

*Polistes* paper wasps are also members of the Vespidae and have smaller, umbrella-shaped paper nests, and colonies of up to several hundred workers, and survive for one year. Mated queens establish nests in spring, and they survive through fall. Typically newly mated queens are the only colony members to overwinter (4). In contrast, solitary aculeate (stinging) wasps are much shorter-lived. Adults consume nectar, lay eggs directly into prey or construct burrows or nests and provision eggs with arthropod prey, and individuals from different generations do not overlap.

In this study, *Polistes* paper wasps and solitary wasps were collected to serve as a comparison to honey wasps. These collections included five species of *Polistes* wasps and members of the solitary wasp families Crabronidae, Ichneumonidae, Mutillidae, Sphecidae, Thynnidae, Pompilidae, and Eumenine wasps from the Vespidae.

#### Sample collection and storage

All insect specimens were collected by net at numerous sites in central and south Texas (Fig. 1, Table S1). Locations were mapped using the R package ggmap (6). Honey wasps were collected from 13 nests in an approximate transect running north/south within the Texan proportion of the *B. mellifica* range, with the exception of five individual honey wasps from the Brackenridge Field Lab in Austin. Polistes, solitary wasps, and bees were collected from flowers by net. Specimens were placed immediately in sterile 15 mL or 50 mL tubes in ice or on cold packs for transportation to the laboratory. All specimens were chilled at 4°C until their guts were dissected on a sterile surface with flame-sterilized forceps. Specimens were dissected within 24 hours of returning from the field and their guts, separated from the sting and venom sacs, were stored whole in 100% ethanol at -80°C. Honey wasp guts were also reserved for bacterial isolation by homogenizing with a sterile pestle in 500µL of 20% (v/v) glycerol and water that had been filter-sterilized (2um Nalgene vacuum filter).

#### DNA extraction, sequencing, and qPCR

DNA was extracted from whole insect guts stored in 100% ethanol using the CTAB/phenol-chloroform extraction method (7). Insect guts were homogenized in a bead beater with approximately 0.5 mL of 0.1mm silica beads in 728 µL of cetrimonium bromide (CTAB) in buffer (Tris-HCL, NaCl, and EDTA), 20 µL of 20 mg/mL proteinase K, and 2 µL of 2-mercaptoethanol, and then incubated at 56 °C overnight. Cellular products were removed from the DNA by adding 750 µL of phenol-chloroform-isoamyl to the gut homogenate, centrifuging at 24,000g for 15 minutes at 4 °C, and removing the upper liquid phase to a new tube. 700 µL of isopropanol and 70 µL 3 M sodium acetate was added and the samples were chilled at 20 °C for 30 minutes for the DNA to precipitate. The samples were centrifuged at full speed (24,000 g) to form a pellet, the supernatant was decanted, then then washed twice in 75% ethanol and 4 °C and then air dried for 30 minutes for remaining ethanol to evaporate. The pellets were resuspended in 50 µL of molecular grade water overnight at 4 °C and then stored in the -20 °C freezer.

The presence of bacteria was confirmed by amplifying the 16S rRNA gene using the universal primers identified by amplifying the 16S rRNA gene the universal primers 27F (5'-AGAGTTTGATCCTGGCTCAG-3') and 1492R (5'- GGTTACCTTGTTACGACTT-3') or the V4 region (515F 5'-GTGCCAGCMGCCGCGGTAA-3’ and 806R 5'-GGACTACHVHHHTWTCTAAT-3’). Each reaction contained 19.875 µL of nuclease-free water, 2.5 µL of Thermopol buffer (New England BioLabs), 0.5 µL of dNTPs, 0.5 µL of each primer, 0.125 µL of Taq DNA polymerase (New England BioLabs), and 1 µL of template DNA. Cycling conditions were as follows: 95°C for 2 min; 35 cycles of 95°C for 20 s, 52°C for 20 s, 68°C for 90 s; then final extension at 68°C for 5 min. Broad range DNA was quantified using a Qubit, DNA samples were diluted to 10 ng/µL and submitted to the GSAF (University of Texas at Austin) for library preparation and sequencing. The V4 region of the 16S rRNA gene was sequenced on an Illumina MiSeq platform amplifying reads of 250bp in length (Hyb515F_rRNA 5'-TCGTCGGCAGCGTCAGATGTGTATAAGAGACAGGTGYCAGCMGCCGCGGTA -3' and Hyb806R_rRNA 3'-TAATCTWTGGGVHCATCAGGGACAGAGAATATGTGTAGAGGCTCGGGTGCTCTG-5').

Quantitative PCR was performed on the DNA extracted from wasp and solitary bee guts to quantify total 16S rRNA gene copies as an estimate of the number of bacteria present per insect. PCRs were conducted on an Eppendorf Mastercycler RealPlex. Each reaction consisted of 5 µL of iTaq universal SYBR Green Supermix (Bio-Rad), 0.5 µL of 3 µM primers (27F 5'-AGAGTTTGATCCTGGCTCAG-3' and 355R 5'-GCTGCCTCCCGTAGGAGT-3'), 3 µL of $H_20$, and 1 µL of DNA template. The cycling conditions consisted of: an initial denaturation step of 95 °C for 3 min, 5 cycles of 95 °C for 5 s, 65-60 °C for 15 s decreasing 1°C per cycle, and 68 °C for 20 s, followed by and 35 cycles of 95 °C for 5 s, 60 °C for 15 s, and 68 °C for 20 s. The standard curves were amplified from target sequence cloned into pGEM-T vector (Promega) and linearized with Apa I. Samples were amplified in triplicate and the mean was calculated for the analysis. The absolute titers of bacterial 16S rRNA gene copies in the specimen guts were plotted using the R package ggplot2 to visualize. Kruskal-Wallis test and Dunn's test for multiple comparisons with Bonferroni adjustment were used to determine statistical difference between *Brachygastra*, *Polistes*, solitary wasps, and native bees.

#### Bacterial isolation and identification

Honey wasp gut homogenates from San Antonio (SA) nests 1 and 7 were grown on HIA (heart infusion agar, BD Difco) with 5% defibrinated sheep's blood or in MRS broth incubated at 35°C in a $CO_2$ incubator set to 5% $CO_2$ or in the anaerobic growth chamber at 31°C. Growth was observed between 1-3 days and the bacteria were identified by amplifying the 16S rRNA gene. PCR reactions consisted of 0.3 µL of Phusion polymerase (NEB), 6 µL of buffer, 0.6 µL of dNTPs (G Bioscience), 1.5 µL each of primers (27F 5'-AGAGTTTGATCCTGGCTCAG-3' and 1492R 5'- GGTTACCTTGTTACGACTT-3'), 30 µL of H2O, and bacterial colonies or 0.5 µL of culture for the DNA template. Colonies were lysed for 10 minutes at 98°C as the initial denaturation step, followed by 35 cycles (98°C for 10 s, 54°C for 20s, and 72°C for 40s). Amplicons were sequenced at the Genome Sequencing and Analysis Facility (UT Austin).

#### Sequence processing and analysis

The default parameters of the Divisive Amplicon Denoising Algorithm 2 (dada2 package version 1.24.0) in R version 4.2.1 were used to process Illumina fastq files (8). Sequences were received from GSAF demultiplexed and with adapters removed. Primers were removed using the trimLeft function, and reads were further trimmed to 230 bases for the forward reads and 240 bases for the reverse reads based on quality profiles. The SILVA 138.2 database was used to assign taxonomy (9). ASVs with mitochondria, chloroplast, or ambiguous assignment were excluded from the analysis as were sequence variants with fewer than two reads in at least two samples using the kOverA function (10). Additionally, we removed 13 ASVs identified as contaminants by the prevalence method of the decontam package (version 1.16.0) using three negative control blank samples and a threshold of 1.0 (11).

The taxonomy assignments for all remaining ASVs were verified by conducting a nucleotide BLASTn of each genetic sequence. We searched the nr/nt nucleotide database of the National Center for Biotechnology Information. We recorded the genus of the top hit and percent sequence identity set to a 98% cutoff (3rd May, 2025), restricting the search to type material with a few exceptions. Uncultured bacteria were used to identify ASVs for *Ca*. Stammerula, *Ca.* Schmidhempelia, and *Wolbachia* since these bacteria are well characterized but do not grow *in vitro*.

Low microbial biomass can present a problem for Illumina sequencing of the V4 rRNA gene. It can lead to the amplification of contaminant DNA, mitochondria and chloroplast DNA, along with cross-contamination during sample preparation and sequencing (12, 13). In the case of gut samples, sequences derived from ingested food may dominate if resident communities are small. To account for these issues, we tracked the number of reads at every filtering step, and samples that lost more than 50% of the reads at one post-dada2 filtering step were removed (table S2). Additionally, samples with fewer than 1500 reads after all filtering steps were removed from the analyses to ensure adequate reads remained to represent the bacterial communities (Table S2). In total, sixteen samples were removed from the dataset: fourteen samples had fewer than 1500 reads remaining at the end of all filtering steps, and one sample lost >80% of reads during decontamination filtering. Notably, this included twelve solitary wasps, two social *Polistes* individuals, and only one honey wasp of the 64 samples included in this study.

Metagenomic sequencing often results in variable read depth AMONG SAMPLES, making it challenging to determine microbial composition. These data can be normalized by subsampling to an equal read depth which can increase the robustness of some statistical tests (14, 15). However, removing read data can lead to incorrect community diversity measures, particularly if rare ASVs are removed (16, 17). The 94 samples included in this study had a 20-fold variation in read number, so the data were subsampled without replacement to 95% of the minimum sampling depth (2,608 reads per sample) and visually confirmed with rarefaction curves (Fig. S1). This resulted in a rarefied data set containing 356 ASVs and 195,985 total reads from 94 insect specimens (Table S4), a reduction from 2,384,753 total reads and 371 ASVs (all removed ASVs had fewer than 10 total reads). To ensure rarefying did not change the outcome of our findings, all statistical analyses were performed on the complete data set and the rarefied data set. We found no difference in the results (tables S5, S6, and S7 ,) The rarefied data were used to present the results in Fig 2, Fig. 3, and Fig. 4.

A heatmap of the Top 50 most abundant ASVs at the genus level was made using ggplot2. Relative abundance plots for the Top 25 and 50 most abundant ASVs at the genus level were created, using the tax_glom function (18) to combine ASVs with the same genus. NMDS plots with Bray-Curtis ordination were made to evaluate the differences in gut bacterial communities between honey wasp nests, and between honey wasps and other wasps. Statistical differences in gut communities between the previously mentioned groups were analyzed using a PERMANOVA with Beta Dispersion Analysis. For the Beta Dispersion Analysis, assumptions were violated; however, the results are still included because populations had natural variation, and the results were statistically significant.

The host-restricted microbiota of honey wasps was compared to *Polistes* social wasps and solitary wasps were identified using the microbiome package core function with prevalence set at 0.5 and detection set at 0 (19). "Core" taxa with a prevalence higher than 0.2 were visualized with heatmaps. Indicator species analysis was performed to determine the bacterial ASVs significantly associated with each honey wasp nest. This was conducted with the indicspecies package, using the multipatt function with statistics calculated with 9999 permutations (20).

#### Phylogenetic reconstruction

The 16S rRNA gene phylogenies were reconstructed for six isolated bacteria and the amplicon sequence variants (ASVs) in high abundance in the honey wasps. Phylogenetic reconstruction was done for sequences in the families Lactobacillaceae, Orbaceae, and Bifidobacteriaceae and they were among the 100 most abundant ASVs following rarefaction. The Bifidobacteriaceae phylogeny included four isolated strains (efB7, efB4, sjhB56, and wjB1), and three ASVs from honey wasps. The Orbaceae phylogeny included two honey wasp ASVs and a bacterial isolate (wjB12). The Lactobacillaceae phylogeny included 16 ASVs isolated from honey wasps and one isolated bacterium (sjhB6). The Lactobacillaceae and Bifidobacteriaceae phylogenies were biased to include 16S rRNA gene sequences from strains isolated from bees and wasps. Sequences of type species representing the major clades of *Bifidobacterium* and genera of the Lactobacillaceae were drawn from GenBank and guided by recent revisions of these taxa (21, 22). Several clades of the Lactobacillaceae phylogeny were collapsed so the full tree showing all branches is included as Fig. S3. The Orbaceae is less well known so the phylogeny included type specimens and sequences from seven uncultured bacteria. Sequences were aligned using the ClustalO in SeaView (23) and manually refined. Maximum likelihood phylogenies were built with the IQ-Tree webserver with 100 standard bootstraps and auto-generated substitution model finder (24). Trees were annotated using the iTol web interface (25).

### Supplementary Figures

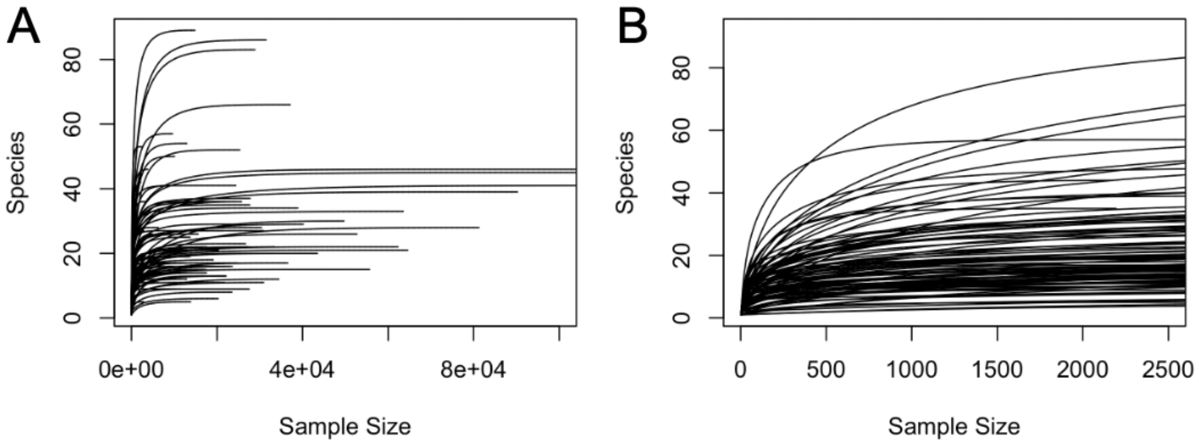


**Fig S1:** Rarefaction curves showing the number of unique species IDs and the sample size (i.e. number of reads) for all insects. (A) shows the total read sample size for all samples, (B) shows the rarefaction curves with the x-axis limited to 2500 reads. All wasp DNA sequences were subsampled to 2063 reads for the rarefied data set (Table S3).

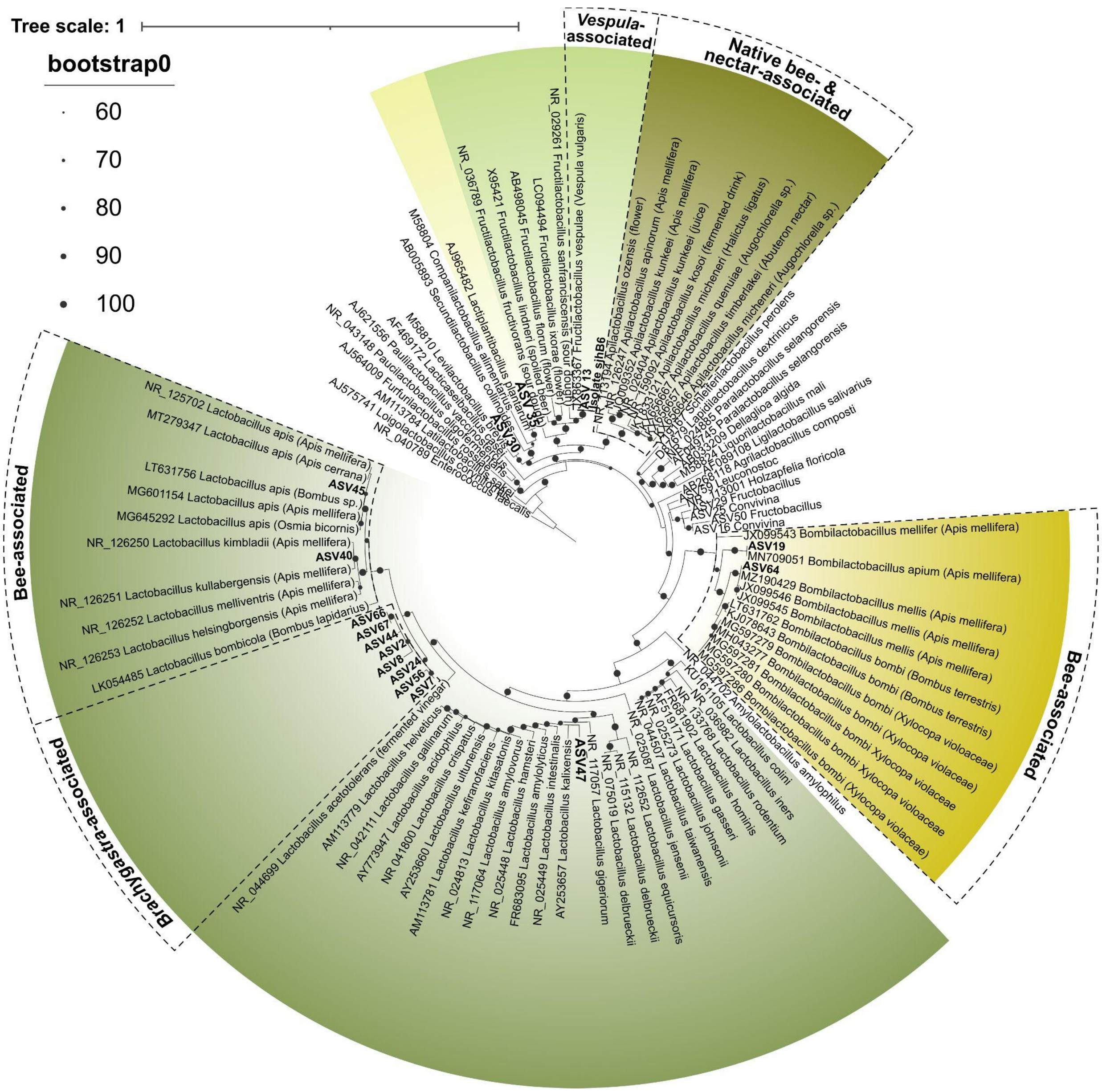


**Fig S2:** 16S rRNA gene Maximum-likelihood phylogeny for Lactobacillaceae*.* The ASVs sequenced from the honey wasps, other wasps, and one honey wasp bacterial isolate have tips highlighted in bold font. Lactobacillaceae representatives are sequences from type specimens retrieved from the NCBI reference database.

#### Statistics Tables

Note: Tables S1-S4 are in a supplementary Excel sheet.

**Table S5**: PERMANOVA test results for gut microbiomes using Bray-Curtis dissimilarity index.

| | | Wasp Groups | | Honey Wasp Nests | |
|---|---|---|---|---|---|
| | | **Full data** | **Rarefied Data** | **Full data** | **Rarefied Data** |
| **Groups** | **Sum of Squares** | 4.669 | 5.488 | 8.628 | 7.8855 |
| | **R-squared** | 0.11952 | 0.14654 | 0.43165 | 0.44258 |
| | **F Statistic** | 12.624 | 15.968 | 2.8627 | 2.9927 |
| | **P Value** | 0.001 (***) | 0.001 (***) | 0.001 (***) | 0.001 (***) |
| **Residual** | **Sum of Squares** | 34.393 | 31.965 | 11.360 | 9.9318 |
| | **R-squared** | 0.88048 | 0.85346 | 0.56835 | 0.55742 |
| **Total** | **Sum of Squares** | 39.062 | 37.453 | 19.988 | 17.8173 |
| | **R-squared** | 1.00000 | 1.00000 | 1.00000 | 1.00000 |

**Table S6:** Homogeneity of dispersion test results for gut microbiomes using Bray-Curtis dissimilarity index.

| | | Insect Groups* | | Honey Wasp Nests | |
|---|---|---|---|---|---|
| | | **Full data** | **Rarefied Data** | **Full data** | **Rarefied Data** |
| **Groups** | **Sum of Squares** | 0.29036 | 0.45756 | 0.68172 | 0.76573 |
| | **Mean Square** | 0.2904 | 0.45756 | 0.02440 | 0.058903 |
| | **F Statistic** | 33.439 | 34.933 | 1.8032 | 1.2641 |
| | **P Value** | 0.001 (***) | 0.001 (***) | 0.063 | 0.253 |
| **Residual** | **Sum of Squares** | 0.80626 | 1.21815 | 1.425 | 2.28322 |
| | **Mean Square** | 0.008669 | 0.01310 | 0.029082 | 0.046596 |

**Table S7:** Paired multiple comparisons between honey wasp nests

| | **Full Data Set** | | | | | **Rarefied Data Set** | | | | |
|---|---|---|---|---|---|---|---|---|---|---|
| **Nest Pairs** | **Sum of Squares** | **F Statistic** | **R-squared** | **P Value** | **Bonferroni adjusted P Value** | **Sum of Squares** | **F Statistic** | **R-squared** | **P Value** | **Bonferroni adjusted P Value** |
| SA1 vs SA2 | 0.2877 | 1.8060 | 0.2051 | 0.1370 | 1.0000 | 0.3016 | 2.2807 | 0.2457 | 0.0840 | 1.0000 |
| SA1 vs SA3 | 0.4597 | 1.8550 | 0.1565 | 0.0590 | 1.0000 | 0.5367 | 2.4417 | 0.1962 | 0.0130 | 1.0000 |
| SA1 vs SA4 | 0.7287 | 3.8829 | 0.2797 | 0.0230 | 1.0000 | 0.7540 | 4.4881 | 0.3098 | 0.0080 | 0.7280 |
| SA1 vs SA5 | 0.3213 | 2.6607 | 0.2754 | 0.0390 | 1.0000 | 0.3170 | 3.3441 | 0.3233 | 0.0140 | 1.0000 |
| SA1 vs SA6 | 1.2412 | 6.7486 | 0.4909 | 0.0150 | 1.0000 | 1.3334 | 9.1667 | 0.5670 | 0.0140 | 1.0000 |
| SA1 vs SA7 | 0.6147 | 4.0476 | 0.3664 | 0.0090 | 0.8190 | 0.5079 | 4.2006 | 0.3750 | 0.0090 | 0.8190 |
| SA1 vs SA8 | 0.8900 | 5.3073 | 0.4312 | 0.0130 | 1.0000 | 0.9863 | 7.6374 | 0.5218 | 0.0120 | 1.0000 |
| SA1 vs AUS | 0.9122 | 3.9200 | 0.3289 | 0.0120 | 1.0000 | 1.0725 | 5.6603 | 0.4144 | 0.0050 | 0.4550 |
| SA1 vs Har1 | 1.1851 | 9.5931 | 0.5159 | 0.0050 | 0.4550 | 1.0338 | 11.7518 | 0.5663 | 0.0040 | 0.3640 |
| SA1 vs Har2 | 0.6043 | 4.6932 | 0.3427 | 0.0030 | 0.2730 | 0.3550 | 4.4693 | 0.3318 | 0.0030 | 0.2730 |
| SA1 vs Individual | 0.9078 | 3.3965 | 0.2740 | 0.0090 | 0.8190 | 1.0509 | 4.6285 | 0.3396 | 0.0060 | 0.5460 |
| SA1 vs KV | 1.4175 | 8.2414 | 0.4518 | 0.0040 | 0.3640 | 1.3101 | 10.2404 | 0.5059 | 0.0030 | 0.2730 |
| SA1 vs PLV | 0.4977 | 2.5508 | 0.2208 | 0.0110 | 1.0000 | 0.4189 | 2.4806 | 0.2161 | 0.0030 | 0.2730 |
| SA2 vs SA3 | 0.2172 | 0.6980 | 0.0907 | 0.6490 | 1.0000 | 0.1740 | 0.5812 | 0.0767 | 0.7570 | 1.0000 |
| SA2 vs SA4 | 0.1895 | 0.8417 | 0.1073 | 0.4660 | 1.0000 | 0.1881 | 0.8345 | 0.1065 | 0.5060 | 1.0000 |
| SA2 vs SA5 | 0.2964 | 2.1763 | 0.3524 | 0.2000 | 1.0000 | 0.2700 | 1.9255 | 0.3249 | 0.3000 | 1.0000 |
| SA2 vs SA6 | 0.5772 | 2.3394 | 0.3690 | 0.1000 | 1.0000 | 0.5925 | 2.5881 | 0.3928 | 0.1000 | 1.0000 |
| SA2 vs SA7 | 0.2566 | 1.3457 | 0.2517 | 0.3000 | 1.0000 | 0.1612 | 0.8669 | 0.1781 | 0.7000 | 1.0000 |
| SA2 vs SA8 | 0.5059 | 2.3169 | 0.3668 | 0.1000 | 1.0000 | 0.5142 | 2.5662 | 0.3908 | 0.1000 | 1.0000 |
| SA2 vs AUS | 0.4495 | 1.4399 | 0.2236 | 0.2070 | 1.0000 | 0.5170 | 1.8293 | 0.2679 | 0.2610 | 1.0000 |

| | Full Data Set | | | | | Rarefied Data Set | | | | |
|---|---|---|---|---|---|---|---|---|---|---|
| **Nest Pairs** | **Sum of Squares** | **F Statistic** | **R-squared** | **P Value** | **Bonferroni adjusted P Value** | **Sum of Squares** | **F Statistic** | **R-squared** | **P Value** | **Bonferroni adjusted P Value** |
| SA2 vs Har1 | 0.4531 | 3.3506 | 0.3583 | 0.0120 | 1.0000 | 0.4014 | 3.4948 | 0.3681 | 0.0130 | 1.0000 |
| SA2 vs Har2 | 0.5490 | 3.8374 | 0.3901 | 0.0170 | 1.0000 | 0.3255 | 3.1891 | 0.3471 | 0.0230 | 1.0000 |
| SA2 vs Individual | 0.6109 | 1.7414 | 0.2249 | 0.1510 | 1.0000 | 0.6841 | 2.1146 | 0.2606 | 0.1150 | 1.0000 |
| SA2 vs KV | 0.6107 | 3.0114 | 0.3008 | 0.0080 | 0.7280 | 0.4060 | 2.4154 | 0.2565 | 0.0390 | 1.0000 |
| SA2 vs PLV | 0.4330 | 1.7851 | 0.2293 | 0.1000 | 1.0000 | 0.3899 | 1.6509 | 0.2158 | 0.2020 | 1.0000 |
| SA3 vs SA4 | 0.1778 | 0.6047 | 0.0570 | 0.7130 | 1.0000 | 0.1668 | 0.5852 | 0.0553 | 0.7160 | 1.0000 |
| SA3 vs SA5 | 0.5117 | 1.8771 | 0.2115 | 0.0800 | 1.0000 | 0.4891 | 1.8673 | 0.2106 | 0.0800 | 1.0000 |
| SA3 vs SA6 | 0.5165 | 1.5382 | 0.1802 | 0.1980 | 1.0000 | 0.5162 | 1.6515 | 0.1909 | 0.1690 | 1.0000 |
| SA3 vs SA7 | 0.3127 | 1.0295 | 0.1282 | 0.4140 | 1.0000 | 0.2001 | 0.6947 | 0.0903 | 0.6780 | 1.0000 |
| SA3 vs SA8 | 0.2722 | 0.8518 | 0.1085 | 0.5350 | 1.0000 | 0.2777 | 0.9375 | 0.1181 | 0.5210 | 1.0000 |
| SA3 vs AUS | 0.4777 | 1.3068 | 0.1404 | 0.2040 | 1.0000 | 0.5702 | 1.6983 | 0.1751 | 0.1200 | 1.0000 |
| SA3 vs Har1 | 0.6007 | 2.4863 | 0.2165 | 0.0240 | 1.0000 | 0.5415 | 2.4844 | 0.2163 | 0.0180 | 1.0000 |
| SA3 vs Har2 | 0.6880 | 2.7874 | 0.2365 | 0.0030 | 0.2730 | 0.6058 | 2.8928 | 0.2432 | 0.0130 | 1.0000 |
| SA3 vs Individual | 0.6479 | 1.6814 | 0.1574 | 0.0590 | 1.0000 | 0.7640 | 2.1397 | 0.1921 | 0.0220 | 1.0000 |
| SA3 vs KV | 0.8117 | 2.9167 | 0.2258 | 0.0150 | 1.0000 | 0.6235 | 2.5458 | 0.2029 | 0.0240 | 1.0000 |
| SA3 vs PLV | 0.5256 | 1.6783 | 0.1572 | 0.1260 | 1.0000 | 0.5181 | 1.7338 | 0.1615 | 0.1210 | 1.0000 |
| SA4 vs SA5 | 0.5968 | 3.1979 | 0.3136 | 0.0310 | 1.0000 | 0.5337 | 2.8402 | 0.2886 | 0.0350 | 1.0000 |
| SA4 vs SA6 | 0.4637 | 1.8563 | 0.2096 | 0.1300 | 1.0000 | 0.4244 | 1.7786 | 0.2026 | 0.1500 | 1.0000 |
| SA4 vs SA7 | 0.3776 | 1.7340 | 0.1985 | 0.1490 | 1.0000 | 0.2687 | 1.2552 | 0.1521 | 0.2910 | 1.0000 |
| SA4 vs SA8 | 0.2719 | 1.1640 | 0.1426 | 0.3050 | 1.0000 | 0.2142 | 0.9639 | 0.1210 | 0.4650 | 1.0000 |
| SA4 vs AUS | 0.5413 | 1.8644 | 0.1890 | 0.0700 | 1.0000 | 0.6356 | 2.3458 | 0.2267 | 0.0390 | 1.0000 |

| | Full Data Set | | | | | Rarefied Data Set | | | | |
|---|---|---|---|---|---|---|---|---|---|---|
| Nest Pairs | Sum of Squares | F Statistic | R-squared | P Value | Bonferroni adjusted P Value | Sum of Squares | F Statistic | R-squared | P Value | Bonferroni adjusted P Value |
| SA4 vs Har1 | 0.3175 | 1.8166 | 0.1679 | 0.1300 | 1.0000 | 0.1857 | 1.1575 | 0.1140 | 0.3160 | 1.0000 |
| SA4 vs Har2 | 0.6903 | 3.8351 | 0.2988 | 0.0070 | 0.6370 | 0.7289 | 4.7993 | 0.3478 | 0.0060 | 0.5460 |
| SA4 vs Individual | 0.9019 | 2.8315 | 0.2393 | 0.0070 | 0.6370 | 0.9746 | 3.2543 | 0.2656 | 0.0070 | 0.6370 |
| SA4 vs KV | 0.7896 | 3.6201 | 0.2658 | 0.0080 | 0.7280 | 0.3857 | 1.9972 | 0.1665 | 0.0600 | 1.0000 |
| SA4 vs PLV | 0.4365 | 1.7721 | 0.1645 | 0.1000 | 1.0000 | 0.4131 | 1.7121 | 0.1598 | 0.1640 | 1.0000 |
| SA5 vs SA6 | 0.9981 | 5.5664 | 0.5819 | 0.1000 | 1.0000 | 1.0343 | 6.3300 | 0.6128 | 0.1000 | 1.0000 |
| SA5 vs SA7 | 0.4549 | 3.6921 | 0.4800 | 0.1000 | 1.0000 | 0.3459 | 2.8724 | 0.4180 | 0.1000 | 1.0000 |
| SA5 vs SA8 | 0.6610 | 4.3802 | 0.5227 | 0.1000 | 1.0000 | 0.6797 | 5.0409 | 0.5576 | 0.1000 | 1.0000 |
| SA5 vs AUS | 0.7173 | 2.7775 | 0.3571 | 0.0370 | 1.0000 | 0.7661 | 3.3276 | 0.3996 | 0.0790 | 1.0000 |
| SA5 vs Har1 | 0.9074 | 10.0534 | 0.6262 | 0.0100 | 0.9100 | 0.6780 | 9.5250 | 0.6135 | 0.0170 | 1.0000 |
| SA5 vs Har2 | 0.5356 | 5.4600 | 0.4764 | 0.0150 | 1.0000 | 0.1728 | 2.9595 | 0.3303 | 0.0180 | 1.0000 |
| SA5 vs Individual | 0.6620 | 2.1643 | 0.2651 | 0.1190 | 1.0000 | 0.7058 | 2.5224 | 0.2960 | 0.0840 | 1.0000 |
| SA5 vs KV | 0.9542 | 5.8095 | 0.4535 | 0.0130 | 1.0000 | 0.8156 | 6.2415 | 0.4714 | 0.0160 | 1.0000 |
| SA5 vs PLV | 0.3974 | 2.0109 | 0.2510 | 0.0900 | 1.0000 | 0.3073 | 1.5961 | 0.2101 | 0.1570 | 1.0000 |
| SA6 vs SA7 | 0.6782 | 2.9009 | 0.4204 | 0.1000 | 1.0000 | 0.5778 | 2.7632 | 0.4086 | 0.1000 | 1.0000 |
| SA6 vs SA8 | 0.3597 | 1.3757 | 0.2559 | 0.4000 | 1.0000 | 0.2088 | 0.9339 | 0.1893 | 0.5000 | 1.0000 |
| SA6 vs AUS | 0.5585 | 1.6108 | 0.2437 | 0.1140 | 1.0000 | 0.6391 | 2.1221 | 0.2980 | 0.1290 | 1.0000 |
| SA6 vs Har1 | 0.3923 | 2.3923 | 0.2851 | 0.0470 | 1.0000 | 0.3421 | 2.6254 | 0.3044 | 0.0350 | 1.0000 |
| SA6 vs Har2 | 1.1481 | 6.6829 | 0.5269 | 0.0190 | 1.0000 | 1.2810 | 10.9011 | 0.6450 | 0.0100 | 0.9100 |
| SA6 vs Individual | 0.8160 | 2.1499 | 0.2638 | 0.0130 | 1.0000 | 0.8934 | 2.6360 | 0.3052 | 0.0170 | 1.0000 |
| SA6 vs KV | 0.8808 | 3.8727 | 0.3562 | 0.0130 | 1.0000 | 0.5055 | 2.7874 | 0.2848 | 0.0530 | 1.0000 |

| | Full Data Set | | | | | Rarefied Data Set | | | | |
|---|---|---|---|---|---|---|---|---|---|---|
| **Nest Pairs** | **Sum of Squares** | **F Statistic** | **R-squared** | **P Value** | **Bonferroni adjusted P Value** | **Sum of Squares** | **F Statistic** | **R-squared** | **P Value** | **Bonferroni adjusted P Value** |
| SA6 vs PLV | 0.7594 | 2.7989 | 0.3181 | 0.0760 | 1.0000 | 0.7801 | 3.1002 | 0.3407 | 0.0720 | 1.0000 |
| SA7 vs SA8 | 0.4307 | 2.0970 | 0.3439 | 0.2000 | 1.0000 | 0.4572 | 2.5321 | 0.3876 | 0.2000 | 1.0000 |
| SA7 vs AUS | 0.4849 | 1.6065 | 0.2432 | 0.1600 | 1.0000 | 0.5011 | 1.8781 | 0.2731 | 0.2340 | 1.0000 |
| SA7 vs Har1 | 0.6556 | 5.1792 | 0.4633 | 0.0150 | 1.0000 | 0.3574 | 3.5159 | 0.3695 | 0.0160 | 1.0000 |
| SA7 vs Har2 | 0.8730 | 6.4954 | 0.5198 | 0.0190 | 1.0000 | 0.3156 | 3.5523 | 0.3719 | 0.0280 | 1.0000 |
| SA7 vs Individual | 0.6985 | 2.0413 | 0.2539 | 0.1210 | 1.0000 | 0.7514 | 2.4217 | 0.2876 | 0.0740 | 1.0000 |
| SA7 vs KV | 0.3358 | 1.7185 | 0.1971 | 0.0800 | 1.0000 | 0.2729 | 1.7404 | 0.1991 | 0.1140 | 1.0000 |
| SA7 vs PLV | 0.6428 | 2.7478 | 0.3141 | 0.0520 | 1.0000 | 0.4099 | 1.8382 | 0.2345 | 0.1170 | 1.0000 |
| SA8 vs AUS | 0.6820 | 2.1051 | 0.2963 | 0.0620 | 1.0000 | 0.7664 | 2.7536 | 0.3551 | 0.0910 | 1.0000 |
| SA8 vs Har1 | 0.6688 | 4.6112 | 0.4346 | 0.0210 | 1.0000 | 0.2758 | 2.4792 | 0.2924 | 0.0470 | 1.0000 |
| SA8 vs Har2 | 0.8989 | 5.8805 | 0.4950 | 0.0250 | 1.0000 | 0.9124 | 9.2657 | 0.6070 | 0.0200 | 1.0000 |
| SA8 vs Individual | 0.8076 | 2.2394 | 0.2718 | 0.0190 | 1.0000 | 0.9246 | 2.8903 | 0.3251 | 0.0150 | 1.0000 |
| SA8 vs KV | 0.5646 | 2.6733 | 0.2764 | 0.0150 | 1.0000 | 0.5585 | 3.3840 | 0.3259 | 0.0110 | 1.0000 |
| SA8 vs PLV | 0.6294 | 2.4939 | 0.2936 | 0.0660 | 1.0000 | 0.5410 | 2.3261 | 0.2794 | 0.0570 | 1.0000 |
| AUS vs Har1 | 0.7278 | 3.2704 | 0.3184 | 0.0070 | 0.6370 | 0.8498 | 4.6494 | 0.3991 | 0.0120 | 1.0000 |
| AUS vs Har2 | 0.9837 | 4.2907 | 0.3800 | 0.0100 | 0.9100 | 1.0410 | 6.0594 | 0.4640 | 0.0080 | 0.7280 |
| AUS vs Individual | 0.6036 | 1.4818 | 0.1747 | 0.0670 | 1.0000 | 0.7796 | 2.1561 | 0.2355 | 0.0170 | 1.0000 |
| AUS vs KV | 0.8134 | 3.0042 | 0.2730 | 0.0080 | 0.7280 | 0.7474 | 3.3837 | 0.2972 | 0.0100 | 0.9100 |
| AUS vs PLV | 0.7141 | 2.2702 | 0.2449 | 0.0340 | 1.0000 | 0.7332 | 2.5568 | 0.2675 | 0.0580 | 1.0000 |
| Har1 vs Har2 | 0.7648 | 7.1555 | 0.4721 | 0.0080 | 0.7280 | 0.8400 | 14.0437 | 0.6371 | 0.0070 | 0.6370 |
| Har1 vs Individual | 1.0954 | 4.1694 | 0.3426 | 0.0050 | 0.4550 | 1.1591 | 5.1314 | 0.3908 | 0.0110 | 1.0000 |

| | Full Data Set | | | | | Rarefied Data Set | | | | |
|---|---|---|---|---|---|---|---|---|---|---|
| **Nest Pairs** | **Sum of Squares** | **F Statistic** | **R-squared** | **P Value** | **Bonferroni adjusted P Value** | **Sum of Squares** | **F Statistic** | **R-squared** | **P Value** | **Bonferroni adjusted P Value** |
| Har1 vs KV | 1.0924 | 6.9416 | 0.4354 | 0.0050 | 0.4550 | 0.2861 | 2.4692 | 0.2153 | 0.0170 | 1.0000 |
| Har1 vs PLV | 0.5624 | 3.0977 | 0.2791 | 0.0350 | 1.0000 | 0.5253 | 3.2750 | 0.2905 | 0.0250 | 1.0000 |
| Har2 vs Individual | 0.9734 | 3.6241 | 0.3118 | 0.0100 | 0.9100 | 1.0101 | 4.6704 | 0.3686 | 0.0270 | 1.0000 |
| Har2 vs KV | 1.4988 | 9.2182 | 0.5060 | 0.0030 | 0.2730 | 1.1091 | 10.3320 | 0.5345 | 0.0050 | 0.4550 |
| Har2 vs PLV | 0.3613 | 1.9277 | 0.1942 | 0.0570 | 1.0000 | 0.3512 | 2.3287 | 0.2255 | 0.0090 | 0.8190 |
| Individual vs KV | 0.9036 | 3.0008 | 0.2501 | 0.0030 | 0.2730 | 1.0326 | 4.0500 | 0.3103 | 0.0020 | 0.1820 |
| Individual vs PLV | 0.6692 | 1.9497 | 0.1960 | 0.0650 | 1.0000 | 0.6940 | 2.1902 | 0.2149 | 0.0670 | 1.0000 |
| KV vs PLV | 1.0678 | 4.6640 | 0.3413 | 0.0040 | 0.3640 | 0.7266 | 3.6927 | 0.2909 | 0.0040 | 0.3640 |

**Table S8:** Summary statistics of 16S rRNA gene copies for different groups of insects

| Insect Group | Sample Size | Median | Mean | Minimum | Maximum |
|---|---|---|---|---|---|
| Honey wasps | 10 | 1.43E+08 | 1.40E+08 | 2.86E+07 | 2.16E+08 |
| Paper wasps | 19 | 3.10E+05 | 2.36E+06 | 5827 | 1.44E+07 |
| Solitary wasps | 30 | 2.78E+04 | 1.52E+09 | 59 | 4.54E+10 |

**Table S9:** Comparison of 16S rRNA gene copies for different groups of insects

<table>
<tr><td rowspan="2">Shapiro-Wilk Test<br>(normality)</td><td>W Statistic</td><td>P Value</td><td colspan="3" rowspan="4"></td></tr>
<tr><td>0.11555</td><td>2.2E-16</td></tr>
<tr><td rowspan="2">Kruskal-Wallis Test</td><td>Chi-squared</td><td>P Value</td></tr>
<tr><td>27.92</td><td>3.878E-06</td></tr>
<tr><td rowspan="4">Dunn’s Test<br>(Multiple comparisons with Bonferroni adjustment)</td><td>Comparison</td><td>Z Statistic</td><td>P Value</td><td>Adjusted P Value</td><td>Level of Significance</td></tr>
<tr><td>Honey wasps vs Paper wasps</td><td>-3.41744</td><td>0.000632</td><td>0.001896</td><td>**</td></tr>
<tr><td>Honey wasps vs Solitary wasps</td><td>-4.99073</td><td>6.02E-07</td><td>1.80E-06</td><td>****</td></tr>
<tr><td>Paper wasps vs Solitary wasps</td><td>-1.66177</td><td>0.096558</td><td>0.289674</td><td>ns</td></tr>
</table>

**Table S10:** Indicator species analysis of the bacterial genera based the rarefied data set

| Honey Wasp Nest | ASV # | Genus | Statistic | P Value | Level of Significance |
|---|---|---|---|---|---|
| Austin Memorial Park | ASV13 | *Fructilactobacillus* | 0.715 | 0.0412 | * |
| | ASV45 | *Bacillus* | 0.695 | 0.0415 | * |
| Harlingen 2 | ASV182 | *Spiroplasma* | 0.622 | 0.0427 | * |
| Austin individual wasps | ASV11 | *Ca.* Stammerula | 0.614 | 0.0436 | * |
| Kingsville | ASV62 | *Lactobacillus* | 0.781 | 0.0328 | * |
| Port Lavaca | ASV43 | *Lactobacillus* | 0.869 | 0.0022 | ** |
| San Antonio 2 | ASV65 | *Lactobacillus* | 0.633 | 0.0461 | * |
| San Antonio 5 | ASV32 | *Secundilactobacillus* | 0.75 | 0.0314 | * |
| San Antonio 6 | ASV25 | *Lactobacillus* | 0.819 | 0.03 | * |
| San Antonio 1, 2 & 5 | ASV44 | Atopobiaceae | 0.691 | 0.0394 | * |
| San Antonio 1, 2 & 3 | ASV30 | *Ca.* Schmidhempelia | 0.639 | 0.0418 | * |
| | ASV14 | *Ca.* Schmidhempelia | 0.636 | 0.0439 | * |
| Harlingen 2, Port Lavaca, San Antonio 1 & 5 | ASV3 | *Bifidobacterium* | 0.686 | 0.0152 | * |
| Harlingen 1, Kingsville, San Antonio 4, 6 & 8 | ASV1 | *Bifidobacterium* | 0.649 | 0.0033 | ** |
| Harlingen 2, Kingsville, San Antonio 2, 5 & 7 | ASV2 | *Lactobacillus* | 0.602 | 0.022 | * |
